# The nonsense-mediated decay RNA-surveillance pathway facilitates the hypoxia response by *C. elegans*

**DOI:** 10.64898/2026.05.06.721854

**Authors:** Calista Diehl, H. Robert Horvitz

## Abstract

Hypoxia (low O_2_) induces physiological, metabolic and behavioral changes. The major hypoxia response pathway is conserved from nematodes to mammals and is driven by activation of the HIF (hypoxia inducible factor) family of transcription factors. Despite a fundamental role of the HIF pathway in regulating cellular homeostasis in response to hypoxic stress, interactions between the HIF pathway and cellular quality-control pathways are largely unknown. Using an unbiased genetic screen, we discovered that in *C. elegans* nonsense-mediated decay (NMD), an evolutionarily conserved mechanism of RNA surveillance, acts in parallel to HIF-1 to selectively enhance specific aspects of the response to HIF-1 activation, possibly through co-regulation of a distinct subset of HIF-1-upregulated genes. Our findings reveal a functional integration between oxygen sensing and RNA surveillance and establish NMD as a key regulator of specific aspects of the HIF-1-driven transcriptional program and physiological response.

## Introduction

Hypoxia is a fundamental physiological stress that drives coordinated changes at the cellular, tissue and organismal levels. In the nematode *C. elegans* this response is regulated by the evolutionarily conserved prolyl hydroxylase EGL-9 (EGLN, PHD, or HIF-PH in mammals) and the transcription factor HIF-1 (hypoxia-inducible factor, HIF-1⍺, HIF-2⍺, also known as EPAS1, and HIF-3⍺ in mammals)^1–3^. In normoxic conditions, EGL-9 uses ambient O_2_ to hydroxylate HIF-1, which is then polyubiquitinated and degraded; by contrast, in hypoxic conditions, EGL-9 lacks its O_2_ substrate and cannot hydroxylate HIF-1, leading to HIF-1 stabilization and activation, which drives adaption to hypoxic stress. In mammals HIF induces critical transcriptional changes affecting cellular metabolism, mitochondrial morphology, angiogenesis, and erythropoiesis^4,5^, making EGLN and HIF crucial for adaptation to hypoxic stress and hypoxia-driven aspects of embryonic development, such as blood vessel formation. EGLN is being explored as a potential therapeutic target for a wide range of disorders, including ischemic stroke, neurodegenerative diseases, and cancer^6–8^. While the core hypoxia response pathway is well defined, if and how additional cellular pathways modulate HIF output to tune the hypoxia response remains poorly understood.

One well-studied mechanism for modulating transcriptional output involves RNA quality-control pathways that regulate transcript stability and integrity. Nonsense-mediated decay (NMD) is an evolutionarily conserved RNA surveillance mechanism that identifies and degrades transcripts with premature termination codons^9^. NMD plays a significant role in maintaining cellular homeostasis ^10–13^, and it is estimated that 1-10% of all cellular transcripts in eukaryotes are either directly or indirectly NMD-regulated^13–15^.

Like other stress responses, hypoxia induces significant shifts in transcription, splicing, metabolism, and general cellular homeostasis. Nonetheless, interactions between the EGL-9/HIF-1 pathway and cellular quality control pathways, such as NMD, are largely unknown. Here we report the discovery that in *C. elegans* the process of NMD is necessary for a complete hypoxia response. Based on the results of a genetic suppressor screen and behavioral and transcriptomic analyses, we conclude that NMD is required for specific HIF-1-dependent behavioral and stress-resistance responses by *C. elegans.* The effects of NMD on the hypoxia response are neither caused by a general downregulation of HIF-1 activity, nor through HIF-1-dependent regulation of NMD, but instead NMD likely acts in parallel to HIF-1 to co-regulate specific aspects of the hypoxia response. Our findings uncover an unexpected integration of oxygen-sensing and RNA-surveillance pathways in *C. elegans* and demonstrate that transcript quality-control mechanisms can actively shape organismal adaptation to hypoxia.

## Results

### *smg-1* function is necessary for complete HIF-1-driven downregulation of egg laying

The activation of HIF-1 in *C. elegans* through either exposure to hypoxia or the loss of EGL-9 function results in numerous physiological changes. For example, HIF-1 activation promotes the retention of eggs inside the uterus, presumably to prevent the deposition of eggs in an unfavorable environment with limited oxygen^16,17^. Animals with this egg-laying abnormal Egl phenotype have a uterus visibly bloated with an abnormally large number of developing eggs (Extended Data Fig. 1A,B) and lay eggs at later embryonic stages than wild-type animals, because eggs have spent longer in the uterus before being laid (Fig. 1A-C). This Egl phenotype of egg retention is completely dependent on a functional *hif-1(+)* gene, as eggs are not retained in *egl-9(-) hif-1(-)* double mutants (Fig. 1D, Extended Data Fig. 1C). Previously we showed that the cytochrome P450 gene *cyp-36A1* functions downstream of *hif-1* to mediate multiple aspects of the hypoxia response, including egg retention^18^. We confirmed that loss of *cyp-36A1* function suppresses the Egl phenotype of *egl-9,* as *cyp-36A1; egl-9* double mutants laid earlier stage embryos than *egl-9* single mutants; however, we further observed that this suppression was incomplete, particularly when animals were grown at the higher temperature of 25°C instead of the more typical 20°C (Fig. 1E). These observations led us to hypothesize that one or more additional genes function with HIF-1 to regulate egg laying independently of *cyp-36A1* activity.

**Figure 1.**
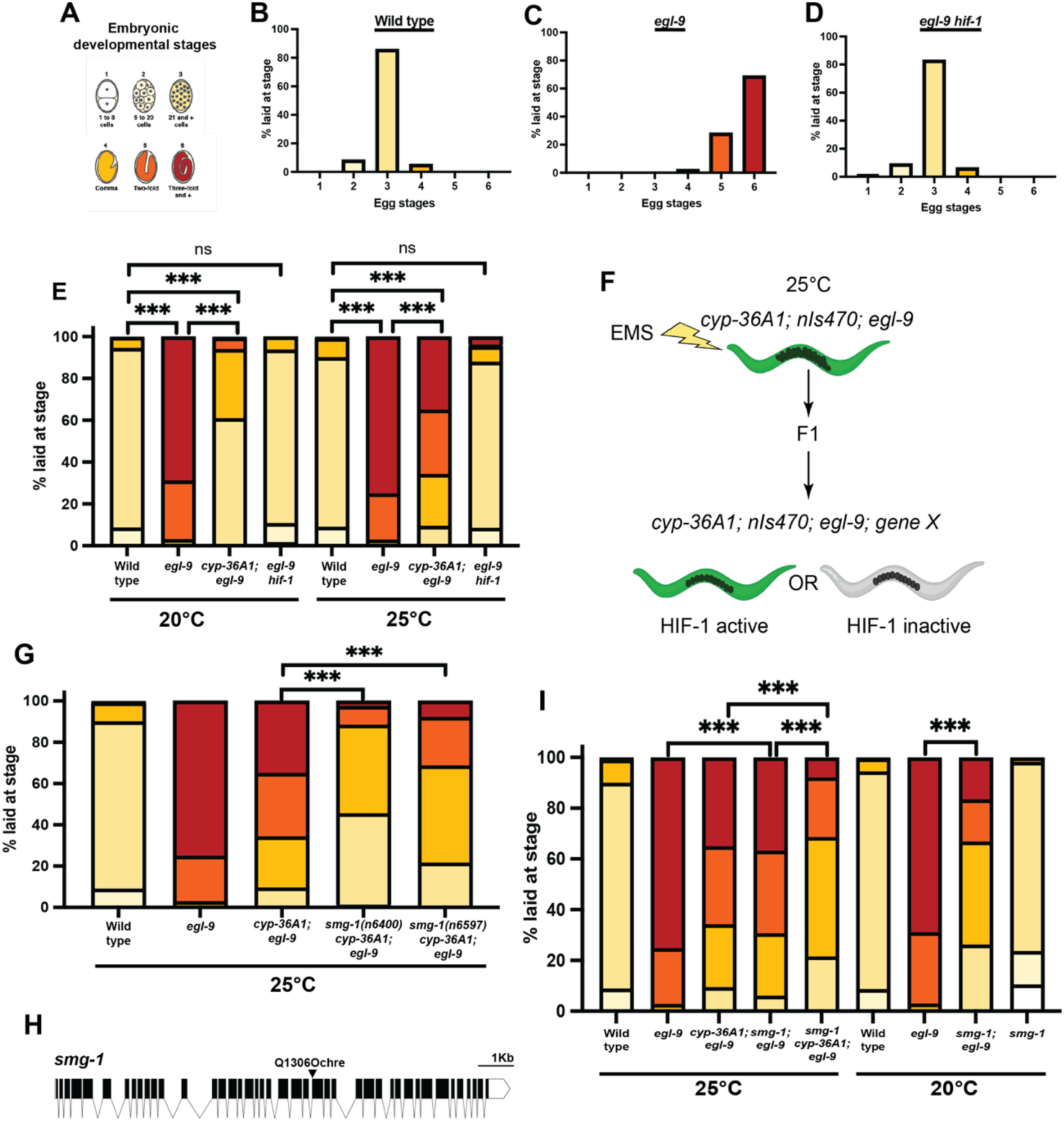
A *smg-1(lf)* mutation suppresses the *egl-9(lf)* Egl phenotype. (A) The six embryonic stages used to categorize recently laid eggs. Sections of bar graphs are matched by color. (B) Wild-type animals lay primarily stage 3 embryos. (C) *egl-9(lf)* mutants lay later stage embryos, producing an Egl phenotype. (D) *egl-9(lf) hif-1(lf)* has wild-type egg laying. (E) *egl-9* Egl phenotype is suppressed by a *cyp-36A1* null mutation at 20^°^C but much less so at 25^°^C. (F) EMS mutagenesis screen to isolate novel *egl-9* suppressors. Strains contain loss-of-function mutations in *egl-9* and the partial suppressor *cyp-36A1. nIs470,* a transcriptional reporter of the HIF-1 target *cysl-2,* is used to indicate HIF-1 activity. (G) Nonsense mutation *n6400* isolated in our screen suppresses *egl-9.* The identical point mutation in *smg-1* generated using CRISPR-Cas9 (*n6597* allele) similarly suppresses *egl-9.* (H) Schematic showing *smg-1* gene structure and the *n6400* and *n6597* Q1306Ochre point mutation in *smg-1.* (I) *smg-1* and *cyp-36A1* function additively. *smg-1(n6597)* suppresses *egl-9* at both 20^°^C and 25^°^C. For all samples N > 100 embryos.

To seek genes that, like *hif-1* and *cyp-36A1,* promote the hypoxia response, we performed an EMS mutagenesis screen for suppressors of *egl-9*. Since a *cyp-36A1* mutation only partially suppressed the *egl-9* Egl phenotype at 25°C, we screened at 25°C for mutations that acted additively with a *cyp-36A1* mutation to further suppress the *egl-9* Egl phenotype to a wild-type non-Egl phenotype. (Fig. 1F). To distinguish those mutations that decreased HIF-1 activity – i.e., either were in *hif-1* itself or that affected factors that acted upstream of HIF-1 –– we included in our mutagenized parental strain *nIs470*, a GFP transcriptional reporter for the gene *cysl-2*, which is expressed in response to HIF-1 activation^19,20^. We sought non-Egl animals in which *nIs470* was expressed, indicating that while the downstream egg-laying behavior was impacted, HIF-1 activity remained intact.

This screen generated an ochre nonsense mutation, *n6400*, in the gene *smg-1* (Fig. 1G,H), which encodes the kinase that activates the process of nonsense-mediated decay (NMD)^21^. Using CRISPR-Cas9 we remade this mutation to generate the *smg-1* allele *n6597* in our starting genetic background and observed a phenotype similar to that of our original isolate, indicating that the nonsense mutation in *smg-1* was the causative mutation (Fig. 1G). Because this *smg-1* allele is a nonsense mutation and causes a recessive phenotype (data not shown) it likely causes a loss of *smg-1* function, indicating that wild-type SMG-1 directly or indirectly facilitates hypoxia-induced egg retention. Our screen isolate maintained expression of the *nIs470 cysl-2* transcriptional reporter, suggesting that *smg-1* is not acting upstream of HIF-1 (Extended Data Fig. 1D-H).

We found that *smg-1; egl-9* double mutants laid earlier stage embryos than *egl-9* single mutants at both 20°C and 25°C (Fig. 1I), indicating that loss of *smg-1* partially suppresses the *egl-9* Egl phenotype independently of *cyp-36A1* function. Additionally, we found that at 25°C *smg-1* and *cyp-36A1* acted additively and thus likely function in parallel rather than in a single linear pathway (Fig. 1I).

### Nonsense-mediated decay enhances the HIF-1-mediated downregulation of egg-laying

s*mg-1* encodes a conserved kinase key in nonsense-mediated decay (NMD), an RNA-surveillance mechanism that identifies and degrades transcripts with premature termination codons. NMD is conserved throughout eukaryotes and is critical for maintaining cellular homeostasis, particularly in response to transcript errors that can result from mutations in DNA or errors in transcriptional or splicing processes^22^. NMD in *C. elegans* involves the functions of seven genes, *smg-*1 through *smg*-7, each of which is necessary for the complete NMD process to occur^23,24^ (Fig. 2A). Unlike in mammals^25^, in *C. elegans* NMD is not necessary for viability, and we observed *C. elegans smg* mutants to have few gross abnormalities compared to wild-type animals. We found that mutations in each of the seven *smg* genes suppressed the *egl-9* Egl phenotype, indicating that the process of NMD is necessary for complete HIF-1 regulation of egg laying (Fig. 2B).

**Figure 2.**
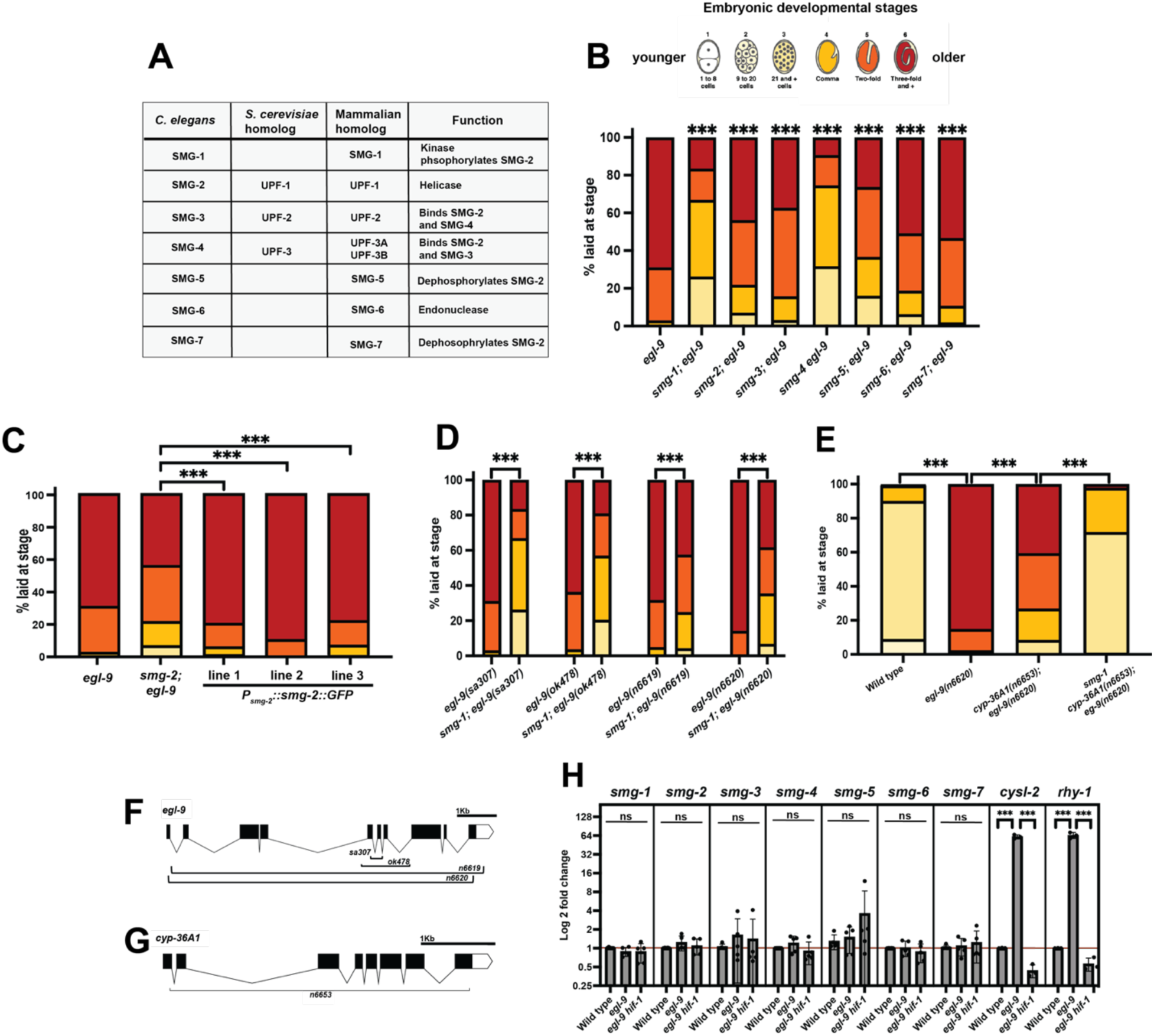
Nonsense-mediated decay enhances HIF-1-mediated downregulation of egg-laying. (A) Table of core NMD proteins and their known functions in *C. elegans*. (B) Loss of any one component of the NMD pathway inhibits the *egl-9* Egl phenotype at 20°C. Statistics show comparison to *egl-9.* (C) *smg-2* suppression of the *egl-9* Egl phenotype is completely rescued by ectopic expression of *smg-2* under its endogenous promoter. (D) *smg-1* suppresses multiple deletion alleles of *egl-9* at 20°C and (E) enhances a deletion of *cyp-36A1* at 25°C. (F) Schematic of the *egl-9* deletions. (G) Schematic of the *cyp-36A1* deletion. For all egg-laying samples N > 100 embryos. (H) qPCR of the seven *smg* genes shows no significant changes among wild type, *egl-9* and *egl-9 hif-1* animals. By comparison, known HIF-1 targets *cysl-2* and *rhy-1* show significant changes. Points represent the triplicate averages of independent biological replicates N = 2-5.

SMG-2 (UPF1 in yeast and mammals) is an RNA helicase core to the NMD pathway. To disrupt the NMD pathway in subsequent experiments, we used both our *smg-1(n6597)* mutation and the well-characterized *smg-2(r863)* allele, which lacks functional NMD^26^. The suppression of *egl-9* by *smg-2* in *smg-2; egl-9* animals was completely rescued by injection of a plasmid containing *smg-2(+)* cDNA driven by the endogenous *smg-2* promoter (Fig. 2C). This result shows that the effects of *smg-2* on egg laying, and likely of other *smg* mutations, are caused by a loss of gene function. Interestingly, similar rescue experiments using tissue-specific promoters (neuron, hypoderm, body-wall muscle, or intestine) indicated that expression in any of these tissues alone was not sufficient to rescue the suppression of *egl-9* by *smg-2* (Extended Data Fig. 2A-D), suggesting that NMD must function in an untested tissue or in multiple tissues to coordinate the EGL-9/HIF-1 egg-laying response.

NMD mutants are often isolated as informational suppressors because of their ability to suppress nonsense alleles^26–28^. We found that *smg-1* was able to suppress multiple distinct *egl-9* alleles, including two deletion mutations (*n6619* and *n6620)* that each removed almost the entire open reading frame (Fig. 2D,F). Additionally, we observed *smg-1* suppression of the *cyp-36A1; egl-9* Egl phenotype at 25°C using almost complete deletion alleles of both *cyp-36A1* and *egl-9* (Fig. 2D-G). These observations demonstrate allele-non-specific suppression of *egl-9* by *smg-1* and establish that NMD is not acting through regulation of *egl-9* or *cyp-36A1* mRNA as an informational suppressor.

Given the role of HIF-1 as a transcription factor, we wondered whether the NMD pathway might be transcriptionally regulated by HIF-1. To quantify mRNA levels of all seven *smg* genes in wild-type, *egl-9* (i.e. HIF-1-activated) and *egl-9 hif-1* (no HIF-1 activation) animals we performed qPCR experiments. We found no significant HIF-1-dependent changes in the expression levels of any of the seven *smg* genes, showing that *smg* transcription is not impacted by HIF-1 activity (Fig. 2H).

Collectively, these data demonstrate that in *C. elegans* loss of NMD reduces HIF-1-driven egg retention, and therefore that NMD activity promotes the HIF-1-driven down-regulation of egg laying.

### Loss of NMD affects some but not all responses to HIF-1 activation

In addition to promoting egg retention, HIF-1 activation causes numerous other physiological and behavioral changes. For example, *egl-9* mutants show decreased rates of locomotion and defecation and increased resistance to various stressors, including hypoxia and oxidative stress^18,29,30^. To determine whether NMD modulates multiple aspects of the hypoxia response, we asked if *smg-1* and *smg-2* mutations impact other HIF-1-dependent responses.

Importantly, we found that the embryonic development and hatching of both *smg-1* and *smg-2* mutants was hypersensitive to hypoxia. Specifically, when grown under hypoxic conditions (0.2% O_2_) approximately 70% of wild-type embryos hatched into first-stage (L1) larvae within 24 hrs, while the remaining 30% arrested in embryogenesis. Consistent with previous studies^29^, we observed that this survival in hypoxia was *hif-1-*dependent, as fewer than 10% of *hif-1* embryos hatched under the same conditions. Both *smg-1* and *smg-2* mutants were significantly more hypoxia-sensitive than the wild type, with only 27-55% of embryos hatching (Fig. 3A). These differences were not simply reflections of baseline changes in developmental timing, as 100% of wild-type, *hif-1*, *smg-1*, and *smg-2* embryos hatched within 24 hrs when grown in normoxic conditions (data not shown). These findings show that NMD interacts with the HIF-1 pathway not only in response to genetic HIF-1 activation in *egl-9* mutants but also in response to environmental hypoxia.

**Figure 3.**
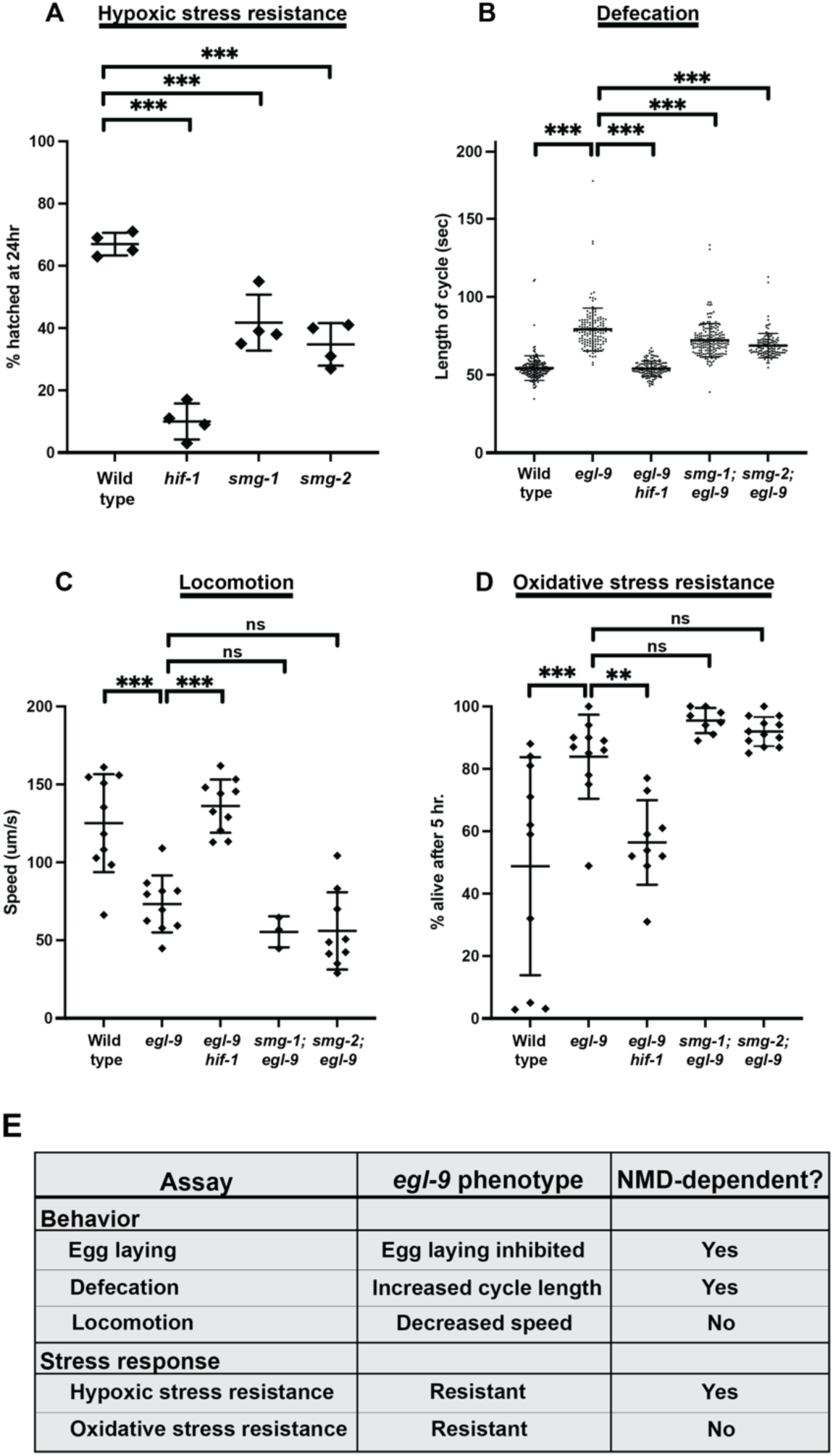
Loss of NMD modulates some, but not all, responses to HIF-1 activation. (A) Both *smg-1* and *smg-2* mutants are hypersensitive to hypoxia compared to wild type, showing more embryonic lethality after 24-hr exposure to 0.2% O_2_. Points show averages of 6 plates each from 4 replicate days. (B) Loss of *smg-1* or *smg-2* mildly suppresses the *egl-9* increase in the defecation motor program cycle length of adults. N > 100 cycles, 18-31 animals. (C) Loss of NMD does not impact the HIF-1-dependent changes in locomotion. Points show plate averages. N = 3-10 plates, 10 animals per plate. (D) Loss of NMD does not impact *egl-9* resistance to oxidative stress, as determined by the percent of adults surviving after 5 hrs on plates containing 7.5 mM t-BOOH. Points show plate averages. N = 8-11 plates, 50 animals per plate. (E) Summary table of all tested HIF-1-depdendent phenotypes.

The *C. elegans* defecation motor program (DMP) involves a coordinated cycle of stereotyped length in actively feeding wild-type animals^31^. Consistent with previous reports^18^, we observed that *egl-9* mutant adults had a decrease in their defecation rate, with each DMP cycle lasting 78 seconds on average, compared to 54 seconds in wild-type adults. Similar to egg laying, this effect on defecation was *hif-1-*dependent (Fig. 3B). This phenotype was also observed for L4 animals, which have not yet begun to develop embryos, showing that the changes to the DMP were not merely a biomechanical consequence of the bloated uterus found in adults (Extended Data Fig. 3A). While loss of NMD alone did not impact adult defecation rates (Extended Data Fig 3B), mutations in either *smg-1* or *smg-2* partially suppressed the *egl-9* DMP phenotype, with average cycle lengths of 71 and 69 seconds respectively (Fig 3B). Thus, similar to the HIF-1-mediated down-regulation of egg laying, the complete HIF-1-mediated down-regulation of defecation requires the function of the NMD pathway.

While loss of NMD impacted multiple aspects of the HIF-1-driven phenotype (reduced egg laying, developmental resistance to hypoxic stress, reduced defecation), we also identified aspects that were NMD-independent. Similar to previous studies, we observed that *egl-9* mutants had a *hif-1-*dependent decrease in locomotion rates^18^. However, the locomotion rate of *egl-9* mutants was not affected by mutations in *smg-1* or *smg-2* (Fig. 3C). Also, previous studies reported that *egl-9* mutants display an increased resistance to oxidative stress as assayed by survival after exposure to the oxidizing agent tert-Butyl hydroperoxide (t-BOOH)^18,30^. We confirmed that *egl-9* mutants displayed increased resistance to t-BOOH and further observed that *egl-9 hif-1* mutants had a wild-type level of t-BOOH sensitivity, showing that this phenotype is *hif-1-*dependent. Unlike resistance to hypoxic stress, the resistance to oxidative stress by *egl-9* mutants was not affected by *smg-1* or *smg-2* mutations (Fig. 3D).

Based on the behavioral and stress-response assays we conducted, we conclude that NMD modulates some but not all aspects of the hypoxia response of *C. elegans* (Fig. 3E). The effects of *smg-1* and *smg-2* mutations on multiple aspects of the hypoxia response –– egg laying, defecation and hypoxic survival –– indicate a substantial integration between the hypoxia and NMD pathways.

### NMD and HIF-1 act in parallel to regulate a specific subset of downstream factors

Given that both HIF-1 and the NMD machinery primarily function to adjust mRNA levels, through transcription and degradation respectively, we hypothesized that the phenotypic differences we observed were caused by changes in mRNA levels. To analyze the effects of both HIF-1 and NMD on mRNA levels we performed whole-animal RNA-Seq studies of young adults of nine genotypes: (1) wild type, (2) *egl-9*, (3) *egl-9 hif-1*, (4) *smg-1*, (5) *smg-1; egl-9*, (6) *smg-1; egl-9 hif-1*, (7) *smg-2*, (8) *smg-2; egl-9*, and (9) *smg-2; egl-9 hif-1* (Fig. 4A). Comparisons of RNAs among these genotypes allowed us to identify transcripts that are regulated by HIF-1 and/or NMD and thereby explore the relationship between NMD and the EGL-9/HIF-1 pathway.

**Figure 4.**
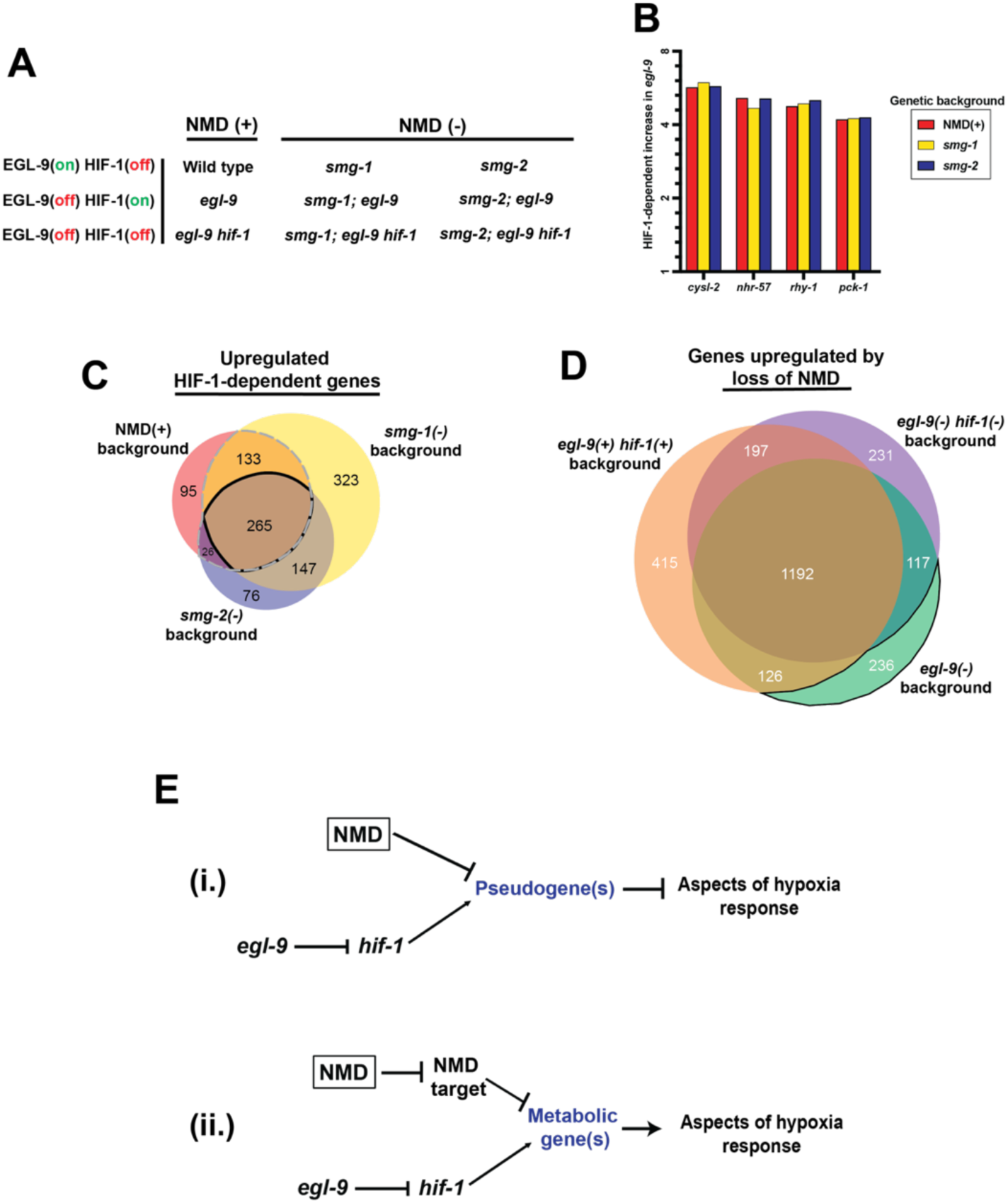
NMD and HIF-1 act in parallel to regulate a specific subset of downstream factors. (A) To help identify possible transcripts impacting the hypoxia response, we performed RNA-sequencing of nine strains with mutations in the EGL-9/HIF-1 and NMD pathways. (B) Fold-change caused by HIF-1 activation in four well-studied HIF-1-dependent genes in the NMD(+), *smg-1(-)* and *smg-2(-)* backgrounds show no significant decrease in HIF-1 activity. (C) Comparisons of genes that are significantly upregulated in a HIF-1-dependent manner in NMD(+), *smg-1(-)* and *smg-2(-)* backgrounds. Dashed line notes the section of genes upregulated in NMD(+) background and at least one NMD(-) background. Solid line notes the section of genes upregulated in all NMD backgrounds. (D) Comparison of genes that are upregulated by loss of NMD in the three different hypoxia backgrounds. Solid line notes the section of genes upregulated in *egl-9(-)* background only. (E) We propose two possible genetic pathways: (i) HIF-1 causes the transcription of a set of aberrant pseudogenes, which are then degraded by NMD, and (ii) both NMD and HIF-1 promote the expression of a subset of metabolic genes. These two models are neither mutually exclusive nor mutually required; they may function separately or together.

To begin, we sought to confirm that loss of NMD did not cause a general decrease in HIF-1 activity, which we did using three criteria. First, we looked at four well-studied HIF-1-upregulated genes: *cysl-2*, *nhr-57*, *rhy-1*, and *pck-1*^32–34^. Corroborating our *cysl-2* reporter results (Extended Data Fig. 1D-H), we found that all four genes were upregulated by HIF-1 to similar levels in NMD(+) and both NMD(-) backgrounds (Fig. 4B). Second, expanding beyond well-studied targets we found that the majority of HIF-1-upregulated genes (increased expression in *egl-9* vs. wild type, and decreased expression in *egl-9 hif-1* vs. *egl-9*; Fold change > 2, Adjusted P-value <0.05) were still differentially regulated in the NMD(-) strains (Fig. 4C, Extended Data Fig. 4A, Extended Data Table 1). Specifically, of the 519 *hif-1*-dependent upregulated genes in the NMD(+) background 82% were also significantly upregulated in at least one NMD(-) background (Fig. 4C dashed outline), and 51% in both (Fig. 4C solid outline). We obtained similar results for HIF-1 downregulated genes (Extended Data Fig. 4B,C, Extended Data Table 2). Third, we examined the magnitude of the HIF-1 upregulation by comparing the increase in *egl-9* vs. wild type in NMD(+) vs. NMD(-) backgrounds. For the 519 *hif-1-*dependent upregulated genes in the NMD(+) background, we found that 64% had similar HIF-1 response levels across NMD(+) and NMD(-) backgrounds –– only 22% had a decrease in HIF-1 magnitude (*smg; egl-9* vs. *smg* <50% of *egl-9* vs. wild type), and only 14% had an increase (*smg; egl-9* vs. *smg* >200% of *egl-9* vs. wild type) (Extended Data Table 3). Collectively, these data establish that loss of NMD does not simply reduce general HIF-1 activity.

Next, we asked if HIF-1 activation causes general increases in NMD activity. While we had already observed that HIF-1 did not cause changes in *smg* transcription levels (Fig. 2H), this finding did not preclude the possibility that HIF-1 might act upstream of NMD through non-transcriptional mechanisms. If HIF-1 caused a general increase in NMD activity then *egl-9* mutants would have increased NMD degradation and thus NMD-target transcript levels would be decreased in *egl-9* mutants. Using a previously published list of candidate *C. elegans* NMD targets^35^, we found that of the 581 putative NMD targets, only 3% showed significant downregulation by HIF-1 activity in the NMD(+) background (Extended Data Table 4). To identify NMD-dependent transcripts in our own data set we compared both NMD(-) strains to the NMD(+) strain of the same *egl-9/hif-1* background, focusing on transcripts that were significantly upregulated in both *smg-1(-)* and *smg-2(-)* backgrounds (Fold change > 2, Adjusted P-value <0.05) (Fig. 4D, Extended Data Fig. 4D, Extended Data Table 5). Of the 2,514 total transcripts that were upregulated after loss of NMD, only 9% were exclusive to the *egl-9* background (Fig. 4D solid outline). Additionally, to determine whether HIF-1 activity impacted the severity of NMD-dependent degradation, we looked at the 1,192 transcripts that were upregulated by loss of NMD in all backgrounds and found that HIF-1 activity increased the magnitude of that effect >2-fold in only 8% of transcripts (Extended Data Table 6). These data show that HIF-1 activity does not cause a general increase in NMD activity.

Since our data indicate that NMD does not act upstream of HIF-1 nor does HIF-1 act upstream of NMD, we proposed a genetic model in which NMD promotes aspects of the hypoxia response in parallel to HIF-1 with the two pathways co-regulating a specific subset of downstream factors.

To identify specific co-regulated transcripts that modulate the hypoxia response we first considered NMD targets with known downstream phenotypic impacts. The mitochondrial tyrosyl-tRNA synthetase gene *yars-2* in *C. elegans*^36^, and multiple components of the integrated stress response (ISR) and unfolded protein response (UPR) pathways in mammals^10,12,37^ (see Extended Data Table 7), have been identified as NMD-targets responsible for regulating stress responses. We examined mRNA levels of genes that encode tRNA synthetases and homologs of mammalian NMD-targeted ISR and UPR pathway components. We found none of these candidate genes displayed both NMD– and HIF-1-dependent regulation (Extended Data Table 7). These results suggest that the integration of NMD and the hypoxia response we observed is likely not mediated by a previously characterized pathway but instead reflects a novel mechanism.

### NMD and HIF-1 oppositely regulate a population of pseudogenes

To identify co-regulated transcripts that might be responsible for controlling the hypoxia response, we first looked at transcripts that were downregulated by NMD and upregulated by HIF-1 activity. Specifically, we first identified transcripts that were significantly upregulated (Fold change > 2, Adjusted P-value <0.05) in *smg-1; egl-9* and *smg-2; egl-9* double mutants compared to *egl-9* single mutants and found 1,855 and 2,104 genes, respectively, with 1,671 genes upregulated in both NMD(-) backgrounds (Extended Data Fig. 4D, Extended Data Table 8). We then looked at those transcripts that were also *hif-1-*upregualted in the NMD(-) background and found 73 transcripts that were both downregulated by NMD and upregulated by HIF-1 activity (Extended Data Fig. 4E, Extended Data Table 8). Using WormCat^38^, a *C. elegans* tool for identifying pathway enrichment, we found that this population of 73 transcripts had an enrichment of pseudogenes, with 16 (22%) of the co-regulated transcripts being pseudogenes (Extended Data Table 8). Based on these data, we suggest a possible model in which HIF-1 promotes the transcription of a set of aberrant pseudogenes that go on to be degraded by NMD; however, in NMD-deficient strains levels of these pseudogenes remain high after HIF-1 activation and they go on to inhibit aspects of the hypoxia response (Fig. 4Ei).

### HIF-1 and NMD both upregulate a set of metabolic genes

While our first analysis looked for co-regulated genes that were upregulated by HIF-1 and downregulated by NMD, we also considered a model in which NMD and HIF-1 both act to upregulate downstream factor(s). This group of genes could be both upregulated directly by HIF-1 and, given the canonical role of NMD in degrading mRNA, upregulated indirectly by NMD. To seek such candidate genes, we identified transcripts that were significantly downregulated (Fold change > 2, Adjusted P-value <0.05) in *smg-1; egl-9* and *smg-2; egl-9* double mutants compared to *egl-9* single mutants and that also showed *hif-1-*dependent upregulation. While there were over 1,000 genes upregulated by loss of NMD, only 69 and 117 genes were downregulated in *smg-1; egl-9* and *smg-2; egl-9,* respectively, with an overlap of 19 genes, only eight of which were also *hif-1*-upregulated (Extended Data Fig. 4F, Extended Data Table 9). Notably, one of these genes was *cyp-36A1,* which has a known role in regulating the hypoxia response^18^; however a *smg-1* mutation suppressed the *egl-9* Egl phenotype even in a *cyp-36A1* complete deletion background (Fig. 2E), precluding *cyp-36A1* from being the sole NMD-dependent regulator of the *hif-1* response.

To get a better sense of possible pathways that were being affected, we relaxed our criteria to include transcripts with a fold change of >1.5. We identified 36 genes upregulated by both NMD and HIF-1 (Extended Data Fig. 4G, Extended Data Table 9), and using WormCat we found that this population had a striking enrichment of metabolic genes, specifically those related to lipid and glutathione metabolism (Extended Data Table 10). Based on these data, we suggest a second, not mutually exclusive, model in which both HIF-1 directly and NMD indirectly promote the expression of a set of metabolic genes that are necessary for a complete hypoxia response (Fig. 4Eii).

## Discussion

We report the discovery of the integration of two fundamental and evolutionarily conserved stress response pathways, the hypoxia response pathway and the RNA-surveillance pathway of nonsense-mediated decay. Specifically, loss of NMD reduced specific aspects of the response to HIF-1 activation (hypoxia resistance, egg retention, and decreased defecation rate), revealing that NMD promotes these aspects of the hypoxia response. We also identified aspects of the response to HIF-1 activation (oxidative stress resistance and decreased locomotion) that were regulated independently of NMD. These findings reveal a previously unrecognized partitioning of HIF-1-dependent physiological responses.

How might NMD and the HIF-1 pathway be integrated to regulate the hypoxia response? We observed neither an NMD-dependent increase in general HIF-1 activity nor a *hif-1-*dependent increase in NMD activity, indicating that neither NMD nor HIF-1 acts strictly upstream of the other. Instead we propose that these two conserved pathways function in parallel to co-regulate one or more downstream genes and pathways. The specificity of this regulation is intriguing, as it suggests that NMD modulation of hypoxic stress is not simply caused by the additive effect of multiple cellular stressors, but instead involves the specific co-regulation of certain phenotypic traits and possibly specific genes. The specificity of this response might help explain why only some HIF-1-dependent responses were impacted by loss of NMD; it is possible that the subset of genes co-regulated by HIF-1 and NMD regulate egg laying, defecation and hypoxic stress resistance, while oxidative stress and locomotion are controlled by other genes not impacted by NMD. The specific tissues and cells in which NMD acts to modulate different aspects of the HIF-1 response are unknown and might provide insights into the specificity of NMD-dependent phenotypes.

Based on our RNA-Seq data set, we identified a population of 16 pseudogenes that are both upregulated by HIF-1 activation and downregulated by NMD. Pseudogenes frequently acquire nonsense and frameshift mutations resulting in premature termination codons that make them common targets of NMD. We suggest that these 16 pseudogenes are direct targets of NMD. We propose a model in which HIF-1 activation promotes the transcription of one or more of these pseudogenes, generating a transcript that is normally degraded by NMD but in animals without NMD is not degraded and instead dampens aspects of the hypoxia response (Fig. 4Ei). Interestingly, eight of these pseudogenes have at least one paralog that was also HIF-1 upregulated, suggesting that HIF-1-upregulation of these pseudogenes might be a byproduct of HIF-1-upregulation of a related functional paralog^39^.

A second population of mRNAs that we explored was those that are upregulated by both NMD and HIF-1 activity, presumably directly by HIF-1 (a transcriptional activator) and indirectly by NMD (which degrades mRNAs). We identified 36 such genes, among which there was significant enrichment for metabolic genes, notably genes involved in lipid and glutathione metabolism. In response to hypoxia, cells undergo significant metabolic reprogramming to prioritize anaerobic processes such as glycolysis, thus making *hif-1*-dependent shifts in metabolism a significant aspect of the hypoxia response across cells types^4,34^. We suggest a model in which the co-regulated metabolic genes are components of this *hif-1*-dependent metabolic reprogramming, and in NMD mutants undegraded NMD-target(s) inhibit the expression of one or more of these metabolic genes, resulting in decreased mRNA levels and a subsequent dampening of the hypoxia response (Fig. 4Eii).

In our models, the possible effects of NMD-targeted pseudogenes and downstream metabolic genes on the hypoxia response are neither mutually exclusive nor mutually required. Either of these two pathways could be sufficient to inhibit the hypoxia response in *smg(-)* strains, or the two might work in parallel. We also note the intriguing possibility that these two models define a single pathway in which in NMD-deficient strains the non-degraded pseudogenes described in Fig. 4Ei are in fact the NMD targets that inhibit the metabolic genes shown in Fig. 4Eii.

Variations in NMD can have profound effects on multiple aspects of human development and disease progression. For example, NMD regulates the splicing and degradation of mRNAs involved in numerous diseases, including β-thalassemia, cystic fibrosis and cancer^40–42^.

Additionally, NMD plays a significant role in neurodevelopment. An estimated 80% of transcripts associated with neurodevelopmental disorders (NDDs) are NMD targets^43^, and variations in multiple components of the NMD complex have been identified as associated with NDDs. Most notably, hemizygous loss-of-function variants in UPF3B^44^, a counterpart of the *C. elegans* protein SMG-4, are among the best established causative variants for NDDs^45–47^. Hypoxia is also a critical factor for NDD, as fetal and perinatal hypoxia exposure have been linked to increased risk of schizophrenia, ADHD and autism spectrum disorder (ASD)^48^, as well as to increased symptom severity among individuals with ASD^49^. Our findings raise the intriguing possibility that humans with NMD-complex variants might have increased sensitivity to hypoxic stress, thereby impacting the incidence and severity of NDDs. This impact might be regulated through increased levels of aberrant pseudogenes and/or through decreases in metabolic pathways such as lipid or glutathione production, the latter of which has been shown to be significantly reduced in individuals with ASD^50^. Our discovery that NMD is necessary for the complete hypoxia response of *C. elegans* reveals a novel integration between oxygen sensing and RNA surveillance. Because the EGLN/HIF and NMD pathways both play critical roles in cellular stress responses during development and disease, this integration might have important implications for understanding how organisms respond to physiological stresses in both normal and pathological settings.

## Acknowledgments

We thank members of the Horvitz laboratory for technical support, advice, and suggestions about the manuscript, and Duanduan Ma and other members of the MIT BioMicro Center for technical support with RNA-Seq experiments. Some strains were provided by the *Caenorhabditis* Genetics Center. We thank the laboratory of Dr. Seung-Jae V. Lee for generously providing all *smg-2* rescue plasmids. Screen schematic was made using images from BioRender.com. This work was supported by the National Institutes of Health (R01GM024663) and the Howard Hughes Medical Institute. H.R.H. is an investigator of the Howard Hughes Medical Institute.

## Author contributions

H.R.H. supervised the project. C.D. and H.R.H. conceptualized the project, designed the experiments and wrote the manuscript. C.D. performed experiments and analyzed data.

## Data and materials availability

The RNA sequencing dataset and subsequent differential expression analysis generated and analyzed during this study is available in the Gene Expression Omnibus (GEO) repository under accession number GSE330000.

## Materials and Methods

### *C. elegans* strains and transgenes

All *C. elegans* strains were cultured as previously described^51^. The N2 Bristol strain was used as the reference wild-type strain. The polymorphic Hawaiian strain CB4856 was used for genetic mapping^52^. All animals were grown at 20°C unless otherwise stated. Transgenes and specific mutations used are listed below:

**LG I:** *cyp-36A1(gk824636, n6653), smg-1(n6400, n6597), smg-2(r863), smg-5(r860)*

**LG III:** *smg-3(r867), smg-6(r896)*

**LG IV:** *nIs470[Pcysl-2::GFP, Pmyo-2::mCherry], smg-7(r1197)*

**LG V:** *egl-9 (sa307, ok478, n6619, n6620), hif-1(ia4), smg-4(ma116)*

### Extrachromosomal arrays

*nEx3309-11[P_smg-2_::smg-2::GFP+P_ges-1_::mCherry], nEx3245-6 [P_dpy-7_::smg-2::GFP+P_ges-1_::mCherry], nEx3248,nEx3330 [P_hlh-1_::smg-2::GFP+P_ges-1_::mCherry], nEx3249-50 [P_ges-1_::smg-2::GFP+P_ges-1_::mCherry], nEx3332-3[P_rgef-1_::smg-2::GFP+P_ges-1_::mCherry]*

### Transgenic strain construction

All *smg-2* rescue plasmids used for transgene construction were generously provided by the laboratory of Dr. Seung-Jae V. Lee^36^. Transgenic strains were generated by microinjection of the germline as previously described^53^. All *smg-2* rescue plasmids were injected at a concentration of 25 ng/μl. The *P_ges-1_::mCherry* plasmid was used as a co-injection marker and was injected at a concentration of either 5 ng/μl (*nEx3245-46, nEx3248-50)* or 10 ng/μl (*nEx3309-11, nEx3330, nEx3332-3)*.

### CRISPR allele construction

CRISPR alleles were generated as previously described^54^. The *n6597* point mutation was generated using a single gRNA and 100 nt ultramer (IDT) containing the desired mutation as the repair template. Deletions *n6653, n6619, n6620* and *n6583* were made using multiple gRNAs throughout the gene of interest and no repair template. For all alleles *dpy-10* was used as a co-CRISPR marker and was outcrossed to the parental strain, removing any *dpy-10* mutation.

### Mutagenesis screen for *egl-9* suppressors

To screen for suppressors of the *egl-9* Egl phenotype that functioned in parallel to *cyp-36A1,* we mutagenized *cyp-36A1(gk824636); nIs470; egl-9(sa307)* animals with ethyl methanesulfonate (EMS) as described previously^51^. Animals were grown at 25°C starting multiple generations before mutagenesis. The starting strain carried *nIs470[Pcysl-2::GFP, Pmyo-2::mCherry],* which contains a transcriptional reporter of the HIF-1-dependent gene *cysl-2,* and served as a readout of HIF-1 activity. This reporter is highly expressed in both *egl-9(lf)* single mutants and our starting strain^19,20^. We used a dissecting microscope to screen the F2 progeny for suppression of the Egl phenotype, picking (1) adults that had fewer embryos in their uterus than *cyp-36A1(gk842636); egl-9(sa307)* mutants, and (2) eggs laid by the F2 animals that were at an earlier developmental stage than those laid by *cyp-36A1(gk842636); egl-9(sa307)* mutants. Screen isolates were backcrossed to determine dominant vs. recessive and single-gene inheritance pattern. The screen allele *n6400*, which conferred a recessive phenotype, mapped to chromosome I based on SNP mapping using a strain containing *cyp-36A1(gk824636); egl-9(sa307)* introgressed into the Hawaiian strain CB4856^52^. Whole-genome sequencing identified a mutation in *smg-1.* Regenerating this mutation with CRISPR demonstrated that it was the causative mutation, as described in the text.

### Egg-laying assay

To quantify egg-laying behavior, we scored the developmental stages of eggs laid by young adult hermaphrodites. Ten L4 animals were picked to a plate one day before scoring. Day-one adults were then moved to a new plate and allowed to lay for 1 hr, after which their progeny were categorized into one of the six developmental stages previously described^55^. Egl-D mutants, such as *egl-9,* retain eggs longer in the uterus, thus laying them at later developmental stages.

### Defecation assay

Ten animals (day-one adults or L4s) were moved to a plate seeded the previous day with 50 μl of *E. coli* OP50 and they were allowed to acclimate for 10 min, after which time the plate was recorded for 10 min at a frame rate of 14 fps using a wormtracker (WormLab). Videos were analyzed and cycles of the DMP counted and time between each expulsion contraction recorded^31^. Data were not included for animals that migrated off food or out of frame for longer than 2 min.

### Locomotion assay

Ten day-one adults were moved to a plate seeded the previous day with 400 μl of *E. coli* OP50 swirled to make a full lawn, and then were allowed to acclimate for 5 min, after which time the worms were recorded for 5 min at a frame rate of 3.5 fps using a wormtracker (WormLab). Videos were then analyzed using the WormLab software to find the average speed for each animal.

### Hypoxia survival assay

The day before the assay NGM plates were seeded with 400 μl of *E. coli* OP50 swirled to cover the whole plate, and 30-40 L4s of each genotype were picked onto a new plate. The following day the full lawn plates were placed under a blower for 30-90 min to fully dry, after which 10 day-one adults were put on each plate and allowed to lay for 1-2 hr before being removed from the plate. Plates were then examined and any embryos that were not developmental stage 3 (either younger or older) were removed. Plates were then placed in a hypoxic chamber (Coy Laboratory) set to 0.2% O_2_. After 24 hr the proportion of L1s and embryos per plate was recorded.

### t-BOOH survival assay

Sensitivity to tert-butyl hydroperoxide (t-BOOH) was assayed by placing 50 day-one adult worms on NGM plates containing 7.5 mM t-BOOH, made using 70% t-BOOH solution (Sigma) and seeded with *E. coli* OP50 bacteria. Survival was evaluated after 5 hrs. Animals that migrated up the side of the plate were not counted.

### RNA sample preparation

Animals were grown on NGM plates, washed off in M9, spun down at 2,000 rpm for 2 min, and excess liquid was removed by aspiration. Samples were then rinsed twice with 3 mL M9 and twice with 3 mL RNase-free water. Excess liquid was removed by aspiration, and the pellet frozen in liquid nitrogen. RLT buffer (QIAGEN) was added to the pellet, and worms were lysed using a BeadBug microtube homogenizer (Sigma) and 0.5 mm zirconium beads (Sigma). RNA was extracted using the RNeasy Mini kit (QIAGEN) according to the manufacturer’s instructions. For qRT-PCR of *smg* genes mixed populations were used. For RNA-Seq 300-500 L4 animals were picked to a new plate and collected 6 hr later as very young adults to avoid significant accumulation of embryos in the uterus.

### qRT-PCR protocol

Reverse transcription was performed using SuperScript IV (Invitrogen). The quantitative PCR with SYBR green dye (Applied Biosystems) was performed by using StepOne Real Time PCR System (Applied Biosystems). Relative quantitation was performed using the ΔΔCt method. *ama-1* or *pmp-3* mRNA level was used as a control for normalization.

### Primers for qRT-PCR

*ama-1* F GGAGCTCGAGTGGATCTTCG

*ama-1* R TTGTGGAGAGTCGGTTGACG

*pmp-3* F ATTGCACATCCCGCATGGA

*pmp-3* R GAGGCGTTTTTGCGACCTTT

*smg-1* F GGGAACCTGATAGAACAGTTTC

*smg-1* R CTCTACATTTTTCGTAGTTTGAAG

*smg-2* F GATTGCTGAGAGCCCGGAGAG

*smg-2* R GATTGCTGAGAGCCCGGAGAG

*smg-3* F CAATCAGCGATGCCTGTAGC

*smg-3* R GCGTGACACGAGTTTGCTTT

*smg-4* F CAAAACCTCCCCGTCCATCT

*smg-4* R TTGGTGGTGACTGAAAGATCGTG

*smg-5* F CGATGTTGCGCAGAAAAGGC

*smg-5* R GATTGAACTTGTACAGTCC

*smg-6* F GTCAAGGCAACGACAGAAGC

*smg-6* R CCGACTCCTTATCGAGCACC

*smg-7* F CAGTCGAAACGGATGAAGATG

*smg-7* R GCATGAAAAGAAAAAAAACCAAAAATTATGAC

*cysl-2* F GAACTTGCACTGGAGTCGGA

*cysl-2* R ATTCCGGTTCCCATGCCTTG

*rhy-1* F GCGAAGTGCGATTGTACTCATC

*rhy-1* R TAAAGTTGCATGTCGAGTCCCA

RNA-Seq library preparation:

Following RNA isolation, RNA integrity and quantity were determined by Fragment Analyzer (Agilent). polyA RNA was isolated from ∼500ng of RNA was prepared using NEBNext poly(A) mRNA Isolation at 1/5 reaction, fragmented for 10 minutes, and prepared for short read sequencing using NEBNext Ultra II Directional RNA Library Prep kit for Illumina at 1/10 reaction with 12 cycles of amplification^56^. Libraries were confirmed by fragment analysis and qPCR and sequenced with 75nt paired end reads on an Element AVITI.

### RNA-Seq data analysis

To ensure that the sequencing reads are of sufficient quality for downstream analyses, quality control (QC) was performed using an in-house pipeline developed by the MIT BioMicro Center. This pipeline includes sequencing error rate estimation, sequencing read complexity estimation, FastQC reports, sample contamination estimation, and fragment size distribution analysis.

Alignment to the WBcel235 reference genome was performed using STAR/2.7.9a, and counting was conducted with RSEM/1.3.0 based on R/3.3.1. Gene-wise counts per sample were merged into a matrix, with each gene in a row and each sample in a column. This matrix serves as the foundation for differential expression analyses. Normalization and differential expression analyses were conducted using DESeq2 to assess gene expression differences across conditions. Sample hierarchical clustering and heat map creation were performed using TIBCO Spotfire 12.4, based on Log2(fpkm+1) values calculated by RSEM/1.3.0. To avoid potential noise from non-coding and non-expressed genes, clustering and heatmap analyses were conducted using only protein-coding genes with expression.

## Statistical analysis

Mann-Whitney U tests were used to compare egg-laying data. Unpaired two-tailed t-tests were used to compare between strains: *smg* mRNA expression as measured via qPCR, survival in hypoxia, defecation rates, locomotion rates, and survival on t-BOOH. In cases of multiple comparisons, a Bonferroni correction was applied. For all graphs: * P < 0.05, ** P<0.01, ***P<0.001

**Extended Data Figure 1.**
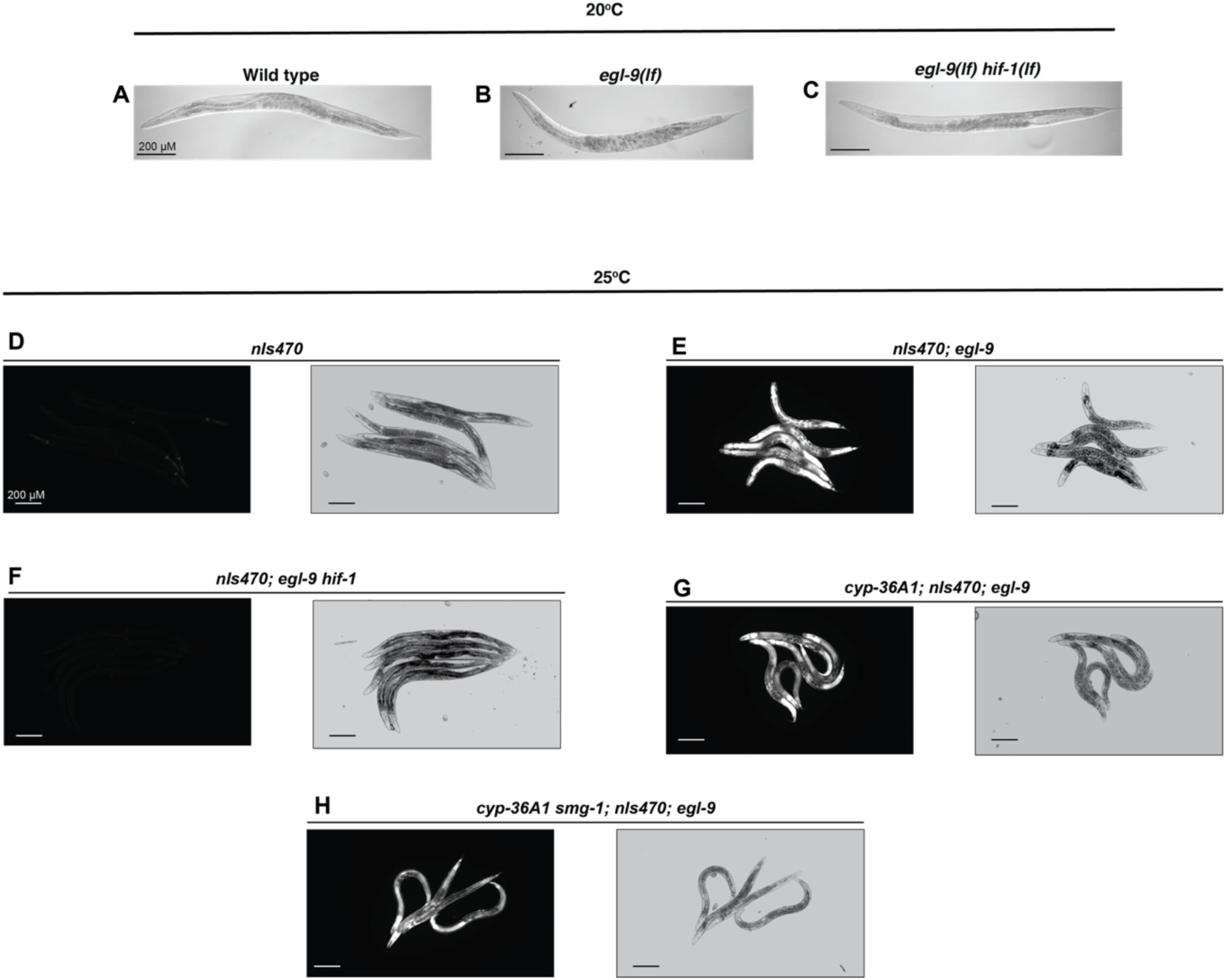
Loss of *cyp-36A1* and *smg-1* do not eliminate HIF-1 activity. Brightfield images of wild type, *egl-9* and *egl-9 hif-1* day-one adults at 20°C. (A) Wild type animals have a single line of embryos in their uterus, (B) while *egl-9* mutants have a uterus bloated with embryos. (C) This effect is *hif-1-*dependent and *egl-9 hif-1* double mutants have uteri similar to wild type. GFP and brightfield images of *nIs470,* a reporter of HIF-1 activity, at 25°C. Reporter expression indicates that HIF-1 activity is low in (D) wild type and (F) *egl-9 hif-1* animals, and high in (E) *egl-9,* (G) *cyp-36A1; egl-9,* and (H) *cyp-36A1 smg-1; egl-9* animals.

**Extended Data Figure 2.**
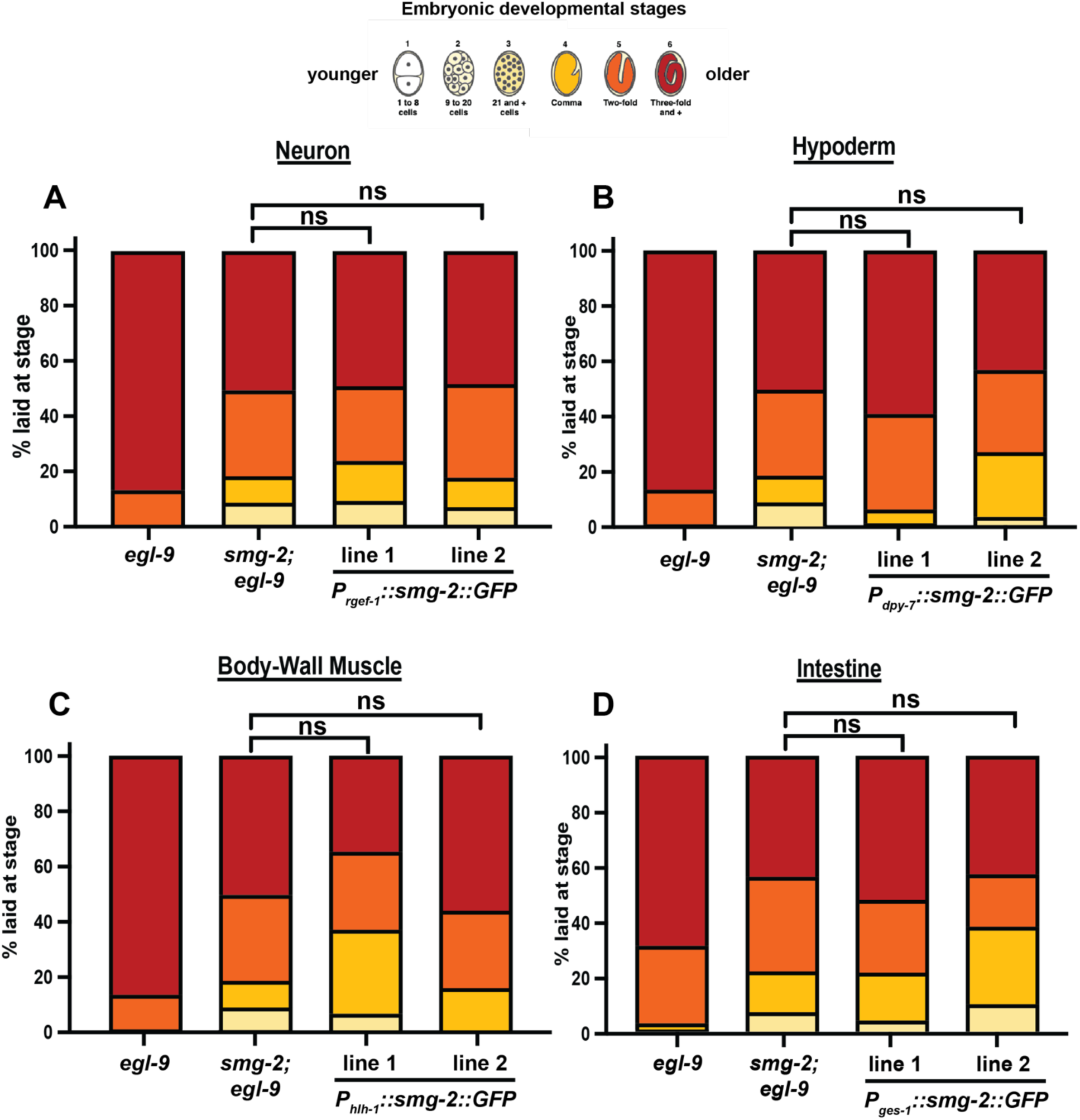
Ectopic expression of *smg-2* in any single tissue is not sufficient to produce the complete hypoxia response. Ectopic expression of *smg-2(+)* under tissue specific promoters in the (A) neurons, (B) hypoderm, (C) body-wall muscle, or (D) intestine is not sufficient to rescue the *smg-2* suppression of *egl-9*. All samples N > 50 embryos.

**Extended Data Figure 3.**
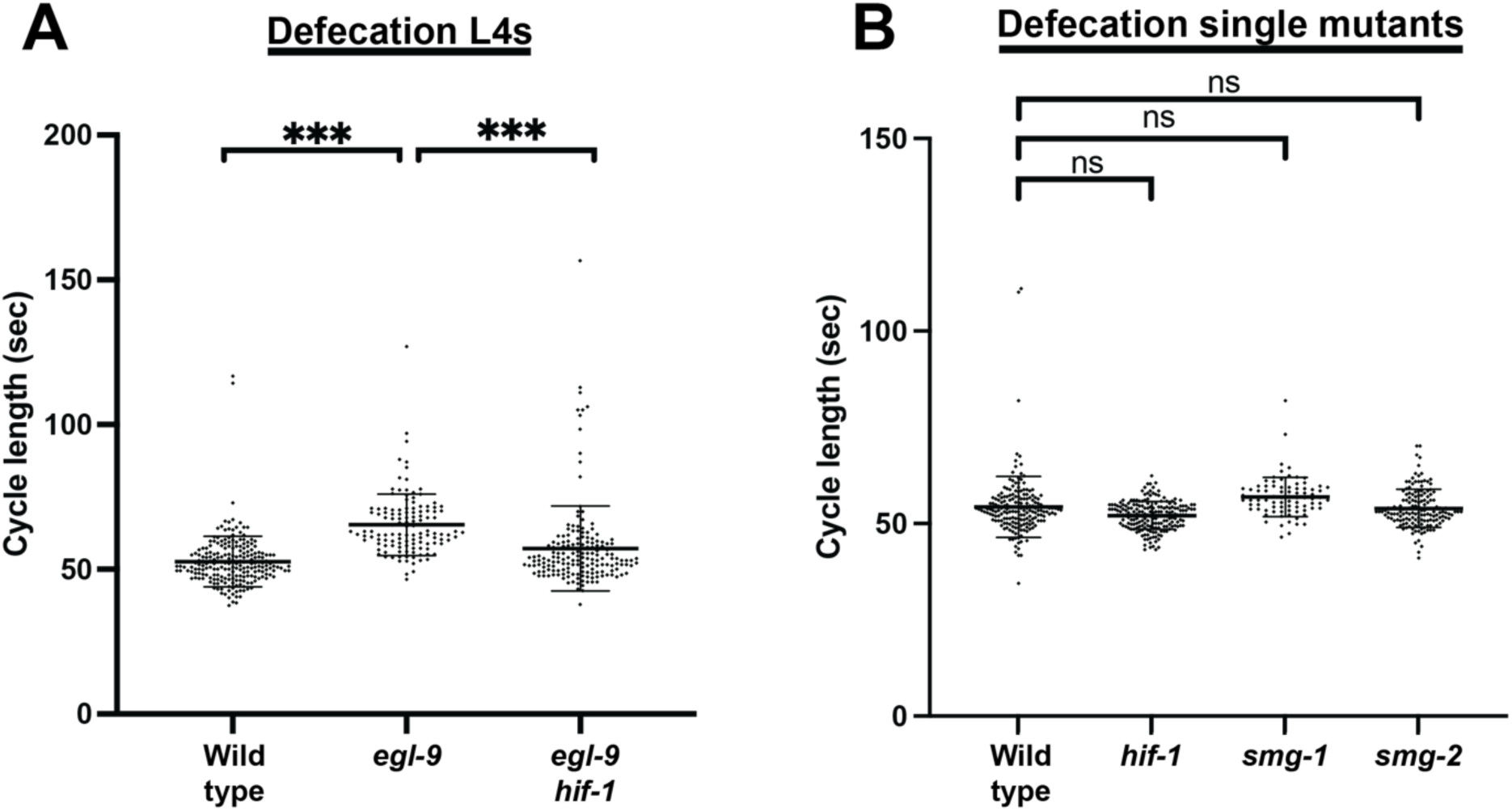
L4 and single mutant defecation. (A) *egl-9* L4s show a decrease in defecation rates similar to that of adults. N > 100 cycles, 17-22 animals. (B) Adult single mutants do not show significant changes in defecation. N > 50 cycles, 9-34 animals.

**Extended Data Figure 4.**
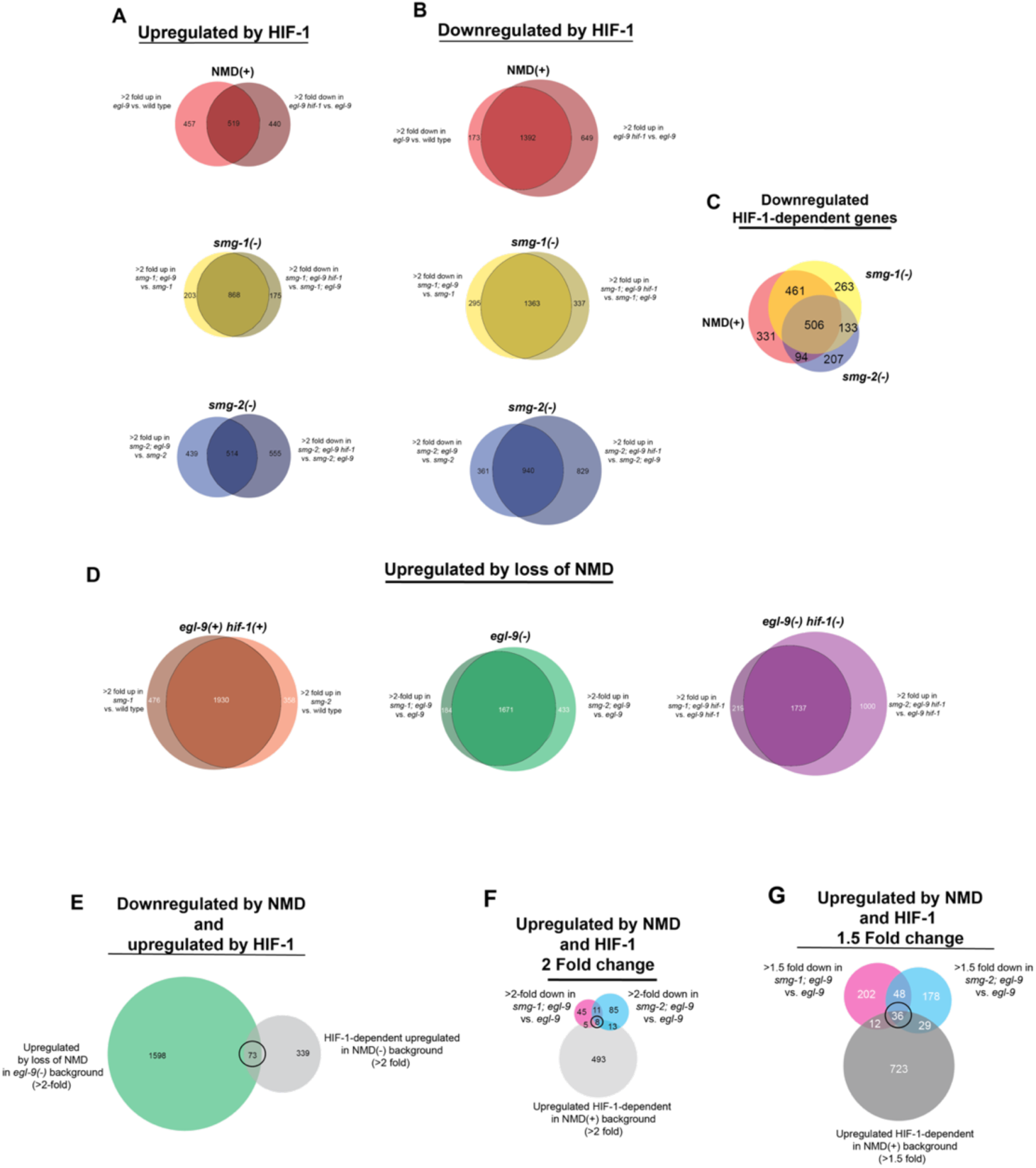
HIF-1 and NMD-dependent transcriptional changes. (A) Genes upregulated in *egl-9* compared to wild type and downregulated in *egl-9 hif-1* compared to *egl-9* in each NMD background (NMD(+), *smg-1(-)* and *smg-2(-)*). Overlapping areas represent *hif-1*-dependent genes shown in Figure 4C. (B) Genes downregulated in *egl-9* compared to wild type and upregulated in *egl-9 hif-1* compared to *egl-9* in each NMD background (NMD(+), *smg-1(-)* and *smg-2(-)*). Overlapping areas represent *hif-1*-dependent genes shown in Figure S4C. (C) Comparisons of genes that are significantly downregulated in a *hif-1*-dependent manner in NMD(+), *smg-1(-)* and *smg-2(-)* backgrounds. (D) Comparison of genes that are upregulated in *smg-1* and *smg-2* vs. *smg(+)* strains in a wild type, *egl-9,* and *egl-9 hif-1* background. Overlapping regions shown in Figure 4D. (E) Comparison of genes that are upregulated in *smg-1; egl-9* and *smg-2; egl-9* vs. *egl-9* (middle region of Figure S4D green *egl-9(-)* Venn diagram), and genes that have *hif-1*-depenendent upregulation in the NMD(-) background (overlap of *smg-1(-)* yellow and *smg-2(-)* blue sections in Figure 4C). Comparison of genes that are downregulated in *smg-1; egl-9* and *smg-2; egl-9* vs. *egl-9* and genes that are upregulated and *hif-1*-depenendent in the NMD(+) background (NMD(+) red section of Figure 4C) based on a (E) >2 fold threshold and (F) >1.5 fold threshold.

## Extended Data Tables 1-10

Table 1: >2-Fold differentially expressed genes up-regulated by HIF-1 activation

Table 2: >2-Fold differentially expressed genes down-regulated by HIF-1 activation

Table 3: Comparison of HIF-1-dependent fold change of HIF-1-upregulated genes between NMD backgrounds.

Table 4: Effect of HIF-1 activation on previously published 581 candidate NMD direct targets.

Table 5: >2-Fold differentially expressed genes up-regulated by loss of NMD

Table 6: Comparison of NMD-dependent fold change of NMD-downregulated genes between *egl-9/hif-1* backgrounds.

Table 7: HIF-1-dependent and NMD-dependent changes in expression of 36 candidate transcripts, 32 tRNA synthetases and 4 homologs of mammalian targets.

Table 8: >2-fold differentially expressed genes up-regulated by HIF-1 activation or down-regulated by NMD.

Table 9: >1.5-fold differentially expressed genes up-regulated by HIF-1 activation or up-regulated by NMD.

Table 10: Functional summary of 36 genes upregulated by both HIF-1 and NMD.

